# Sciviewer enables interactive visual interrogation of single-cell RNA-Seq data from the Python programming environment

**DOI:** 10.1101/2021.08.12.455997

**Authors:** Dylan Kotliar, Andres Colubri

## Abstract

**Summary:** Visualizing two-dimensional (2D) embeddings (e.g. UMAP or tSNE) is a key step in interrogating single-cell RNA sequencing (scRNA-Seq) data. Subsequently, users typically iterate between programmatic analyses (e.g. clustering and differential expression) and visual exploration (e.g. coloring cells by interesting features) to uncover biological signals in the data. Interactive tools exist to facilitate visual exploration of embeddings such as performing differential expression on user-selected cells. However, the practical utility of these tools is limited because they don’t support rapid movement of data and results to and from the programming environments where the bulk of data analysis takes place, interrupting the iterative process. Here, we present the Single-cell Interactive Viewer (Sciviewer), a tool that overcomes this limitation by allowing interactive visual interrogation of embeddings from within Python. Beyond differential expression analysis of user-selected cells, Sciviewer implements a novel method to identify genes varying locally along any user-specified direction on the embedding. Sciviewer enables rapid and flexible iteration between interactive and programmatic modes of scRNA-Seq exploration, illustrating a useful approach for analyzing high-dimensional data.

**Availability and implementation:** Code and examples are provided at https://github.com/colabobio/sciviewer

## 1. Introduction

Dimensionality reduction methods such as UMAP (Becht et al. 2018) and tSNE (Amir et al. 2013) create 2D representations of scRNA-Seq data that preserve distances between cells, providing a visualization that captures much of the underlying data structure. scRNA-Seq analysis can be thought of as identifying, characterizing, and interpreting the biological signals that give rise to that structure. Software to aide in this task includes programmatic toolkits such as Scanpy (Wolf, Angerer, and Theis 2018) and SEURAT (Stuart et al. 2019) for Python and R respectively, and interactive viewers such as Single Cell Explorer (Feng et al. 2019) and CellXGene VIP (Li et al. 2020). While programmatic toolkits provide flexible commands for preprocessing, statistical analysis, and plotting of scRNA-Seq data, they interface with the user solely via programming commands and don’t allow visual interaction with the embedding (e.g. selecting cells). Interactive interfaces enable direct visual interaction but generally do not support the flexibility of the programmatic toolkit. To our knowledge, no interactive scRNA-Seq visualization tool currently supports real-time transfer of data and results to and from the programming environment. Such transferability could enable users to rapidly iterate between interactive discovery of visual patterns, and computational analysis to validate those patterns. We therefore developed Sciviewer to facilitate interactive visual exploration of 2D embedding from within the Python programming environment.

## 2. Methods

Sciviewer is implemented with the Processing data visualization API in Java (https://processing.org/) which is accessible from within Python via the Py5 package (http://py5.ixora.io/about/). We leverage the hardware-accelerated rendering engine in Processing (Colubri and Fry, 2012), which can handle complex geometries in real time, to visualize large scRNA-seq datasets during interactive manipulation. It requires two inputs: (1) a gene expression matrix **X** (N cells X G genes, *X*_*i,g*_ denotes expression of gene *g* in cell *i*), and any 2D embedding of the data such as UMAP - **E** (N cells X 2 dimensions, (*E*_*i,x*_, *E*_*i,y*_) denotes coordinates for cell*i*).

Sciviewer is launched from Python, and opens as a graphical interface that includes an interactive scatter plot of the embedding (Figure 1). Users can select a group of cells {*i*_1_… *i*_*k*_} to compute differential expression between selected and unselected cells. Sciviewer then shows the list of the most differentially expressed genes (defined via Welch’s T-test), alongside violin plots of user-selected genes (Figure 1C). Alternatively, Sciviewer can identify genes that vary locally along any direction in the embedding (Figure 1B). Users select a set of cells and a direction*v* = (*v*_*x*_, *v*_*y*_) and Sciviewer calculates the vector projection of the selected cells onto that direction and displays the genes with the greatest Pearson correlation (*R*_*g*_) between the projected coordinates and gene expression. Mathematically, for gene *g*:

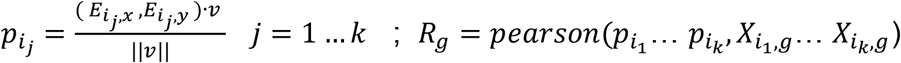

**Figure 1.**
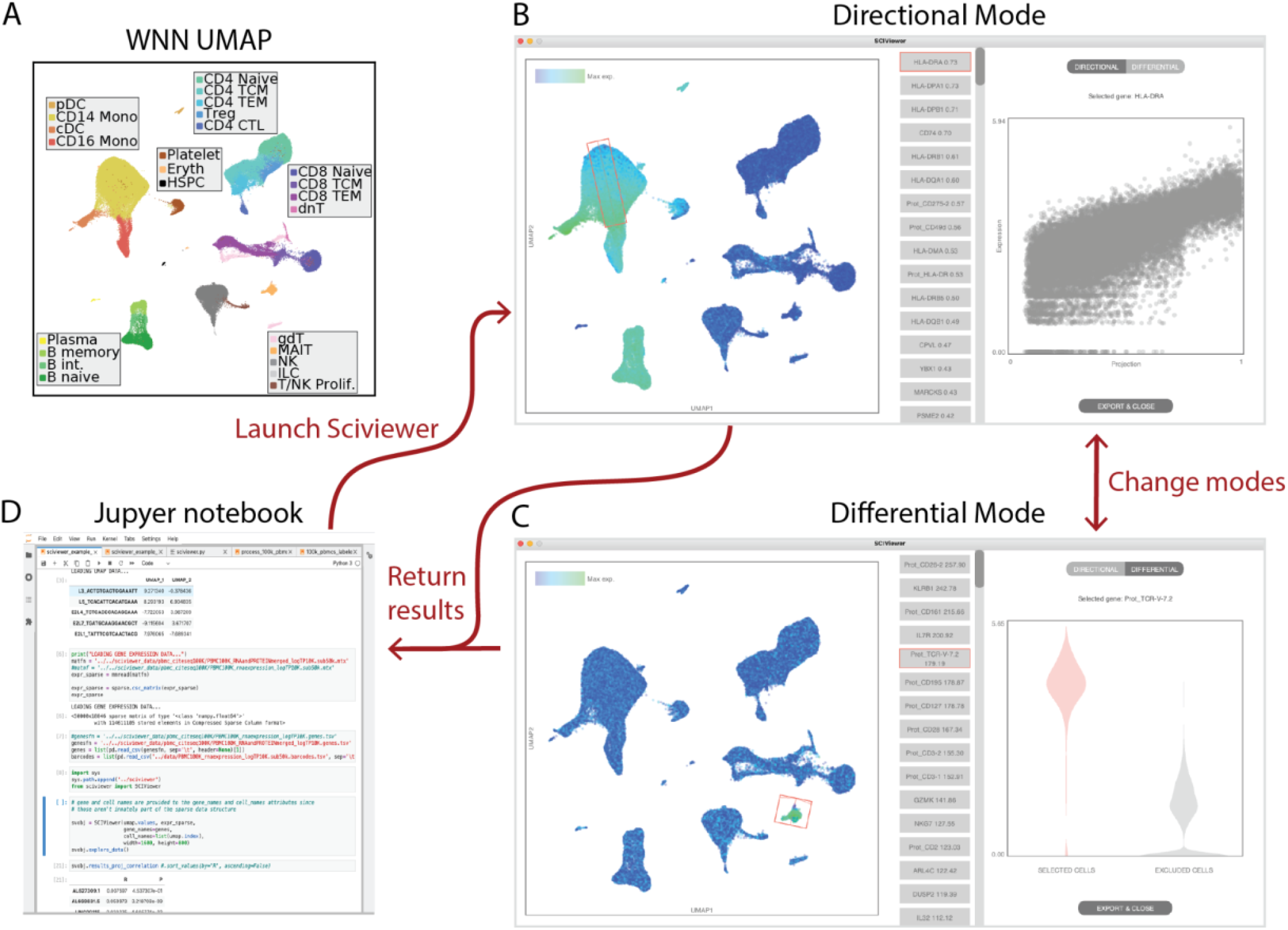
Application of Sciviewer to multimodal PBMC dataset. (**A**) UMAP embedding of CITE-Seq data of peripheral blood mononuclear cells (PBMCs) described in (Hao et al. 2021). Cells are labeled based on clusters described in that paper, with related cell-types aggregated for ease of visualization. (**B-C**) Screenshots of Sciviewer in directional and differential mode respectively for different user selections. (**D**) Screenshot of a Jupyter notebook environment, from which Sciviewer is called, and in which results of Sciviewer selections and calculations are available for programmatic analysis in real time.

This is analogous to pseudotemporal ordering (Saelens et al. 2019), but the ordering is defined by a user-selected direction, allowing for rapid and flexible interrogation of the embedding. Notably, actions in Sciviewer cause real-time updates to corresponding variables in Python so users can programmatically access the selected cells, and associated genes, test statistics, and P-values, for downstream programmatic analyses such as gene-set enrichment (Figure 1D).

## 3. Results

To illustrate the insights obtainable with Sciviewer, we applied it to a CITE-Seq dataset of 161,764 circulating immune cells, and 17,516 genes, consisting of transcriptome-wide profiling and targeted antibody-based capture of 211 proteins (Hao et al. 2021). We use Sciviewer to explore the novel weighted nearest neighbor-based UMAP described in the paper (Figure 1A), which intelligently weights protein and RNA data to generate the embedding. This demonstrates how Sciviewer is agnostic to the choice of 2D embedding and allows us to characterize signal from both RNA and protein features. Directional analysis of CD14+ monocytes demonstrated a gradient in expression of multiple HLA genes (responsible for antigen presentation) at both the RNA and protein levels, thus connecting a biological signal to the organization of the embedding (Figure 1B). Selecting a cluster of cells labeled as mucosal associated invariant T-cells (MAITs) in directional mode, we note a T-cell receptor V-segment protein that is not associated with any of the other T-cell populations, indicating the “invariant” receptor aspect of this T-cell population (Figure 1C). On a 3.8 GHz 8-core Intel Core i7 Mac desktop computer, for this large dataset, it took 3.69 seconds to compute directional correlations for a selection of 25,531 cells, and 7.3 seconds to compute differential expression for 20,661 cells compared against 141,103 others, demonstrating the performance of the tool for a large dataset. This dataset and others are available as part of Sciviewer tutorials in the Github repository.

In summary, Sciviewer enables interactive exploration of scRNA-Seq that is tightly integrated with programmatic analysis in Python. It also introduces a novel directional association analysis that enables flexible exploration and interpretation of 2D embeddings. This approach could potentially have broad utility for other high-dimensional data types beyond scRNA-Seq.

## Funding

The project described was supported by award Number T32GM007753 from the National Institute of General Medical Sciences. The content is solely the responsibility of the authors and does not necessarily represent the official views of the National Institute of General Medical Sciences or the National Institutes of Health.

## Notes

https://github.com/colabobio/sciviewer

### Competing Interest Statement

The authors have declared no competing interest.

